# Cooperativity through Allee Effects drives growth of diffuse midline gliomas impacting the optimal scheduling of therapeutic interventions

**DOI:** 10.64898/2026.09.25.753722

**Authors:** Haider Tari, Ketty Kessler, Elisa Izquierdo, Andrea Sottoriva, Chris Jones

**Affiliations:** Centre for Children and Young People’s Cancer, Division of Cancer Biology, Institute of Cancer Research, London; Institute of Medical and Molecular Genetics (INGEMM), La Paz University Hospital, Madrid, Spain; Computational Biology Research Centre, Human Technopole, Milan, Italy

## Abstract

Diffuse midline gliomas are a group of tumours for which no effective therapies exist. These tumours have substantial intratumoural heterogeneity and harbour beneficial subclonal interactions. Therapeutic failure due to resistance is a major concern in these tumours. We demonstrate how distinct subpopulations derived from the same tumour can interact with one another to alleviate the impacts of therapeutic intervention. This is achieved through the establishment of a mathematical extension to the Lotka-Volterra competition model, where interactions perturbing the Allee effect were demonstrated to exist. These interactions had substantial downstream consequences as they were determined to be perturbed by therapeutic interventions and strikingly, when simulating a two-drug paradigm, the interactions were demonstrated to substantially later the outcomes of treatment scheduling.

## Introduction

Intra-tumoural heterogeneity (ITH) is a commonly observed facet of tumours, which confounds our understanding of cancers, ultimately leading to poorer patient prognosis [1–5]. Paediatric-type diffuse high-grade gliomas (PDHGG) are a highly heterogeneous group of tumours with no effective therapeutic options and little improvement in clinical outcome for decades [6–9]. Diffuse midline gliomas (DMG) typically occur in young children, and originate in critical regions within the brain, such as the brainstem, complicating surgical interventions [10–12]. These tumours display a low mutational burden but a high degree of ITH [13], coupled with key somatic driving mutations leading to the arrest of neurodevelopmental differentiation programs, in common with many paediatric malignancies [13,15,16].

Beyond the diverse landscape ITH represents, it has been demonstrated that the interactions present between different subclonal populations can be a driver of tumorigenic phenotypes [7,17,18] These interactions, between subclonal populations and with their environment, have been linked to an enhanced ability of tumours to grow, disseminate, evade the immune system, and lessen the impact of therapeutic interventions [1,19–23].

The added complexity presented by subclonal interactions means that they are often difficult to study and measure, requiring a shift in both experimental and analysis paradigms to focus on their effects. However, these interactions can be studied through the lens of evolution and ecology, with the aim of describing the altering dynamics created by the presence of multiple distinct interacting subclonal populations [24]. In ecological theory, interactions between two populations can be beneficial, detrimental or have no effect, and these are bidirectional but not strictly symmetric, with both populations affecting one another in potentially distinct modes. [24,25]. These interactions are frequently observed within cancer, with Darwinian evolution explaining the ability of a single clone to perform a selective sweep, dominating the tumour ecosystems. However, other interactions, specifically those conferring a benefit are less frequently observed due to instability of cooperation between populations that occupy the same niche within a system [26–28]. This means that whilst interactions could exist, they do so within an evolving landscape and therefore are not guaranteed to be present when observed. Nonetheless, several studies have demonstrated and measured the effect of these interactions, with exploration across a range of tumour types, such as breast, prostate, colorectal, lung and PDHGG [3,7,17–20,28,29].

Although these interactions have been demonstrated across a landscape of tumours, the precise quantification of their effects can be difficult to achieve. Here, mathematical and computational modelling have been frequently employed to infer the ecological dynamics present [30]. When applied to subclonal interactions they can provide strong analytical and descriptive tools to develop our understanding [18,31,32].

This study aims to explore how subclonal interactions present in the early stages of cell culture growth can modulate the conditions to promote enhanced cell growth.

Specifically, we focus on the Allee effect, which occurs in populations at low densities [23], and how subclonal interactions can lessen its impact.

A pair of primary DMG cell lines derived from a single patient at rapid autopsy are utilised as our experimental system. These cell lines comprise a parental culture (unexposed to therapeutic selection) and a resistant line derived through long-term therapeutic selection with a clinically approved agent, trametinib. Through inference powered using mathematical modelling, we quantify the underlying interaction dynamics present between these cell lines and are able to illuminate the implications for tumour dynamics.

## Results

### Coupled experimental and mathematical approach to quantify subclonal interactions

The experimental design in this study involves culturing two distinct cell lines derived from the same patient tumour. From B169-Parental (referred to as B169 henceforth), we have previously derived B169-T3 (referred to as T3 henceforth), a trametinib-resistant line established via cell culture under continuous trametinib exposure (see Material and Methods). [33]

We grow both lines in isolation and in 3 co-cultures where we seed B169 as 25%, 50% or 75% of the population with the remainder being made up by seeding T3. These cells were imaged at 6 hours intervals, with fluorescent labels allowing for the identification of each population.

Within our experimental approach we used confluence as a metric of growth instead of total cell number. This is because it captures the most challenging phenotype of PDHGGs, their diffuse growth behaviour. PDHGGs are known to invade beyond the initial tumour site, spreading to critical regions in the brain. Measuring confluence as our metric considers both the total cell numbers as well as their diffusivity.

### B169 and T3 glioma lines display markedly distinct growth dynamics

To illuminate potential interactions between distinct populations, we begin by quantifying the phenotype of each population individually. We achieve this by employing ordinary differential equations (ODE), which in this case describes how the population confluence changes with respect to time.

We explored the suitability of several growth models, including the common choices of exponential and logistic growth models. However, as we will demonstrate, the most appropriate model was a weak-Allee effect model, which is an extension of the Allee effect growth model [24]. The Allee effect in general describes a positive relationship between the per-capita growth rate while the population size is relatively small. The weak Allee effect has been demonstrated to be relevant to cancer, specifically in describing *in vitro* systems. This is not a surprising phenomenon as it is widely known that cell-cell communication, for example by chemokine signalling, are crucial to optimal growth. The weak Allee effect can be described using the following ODE:

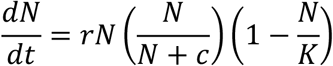

where the new term c represents the Allee effect threshold. Under this model the growth of a population is slower initially, while the population is a smaller size.

To identify the most appropriate model, we employed the use probabilistic Bayesian inference (see Methods) to fit the ODE models to our experimental data. Among the range of models explored, the Weak-Allee effect model had the lowest LOOIC (leave-one-out cross-validation information criteria) (Figure 1A).

**Figure 1:**
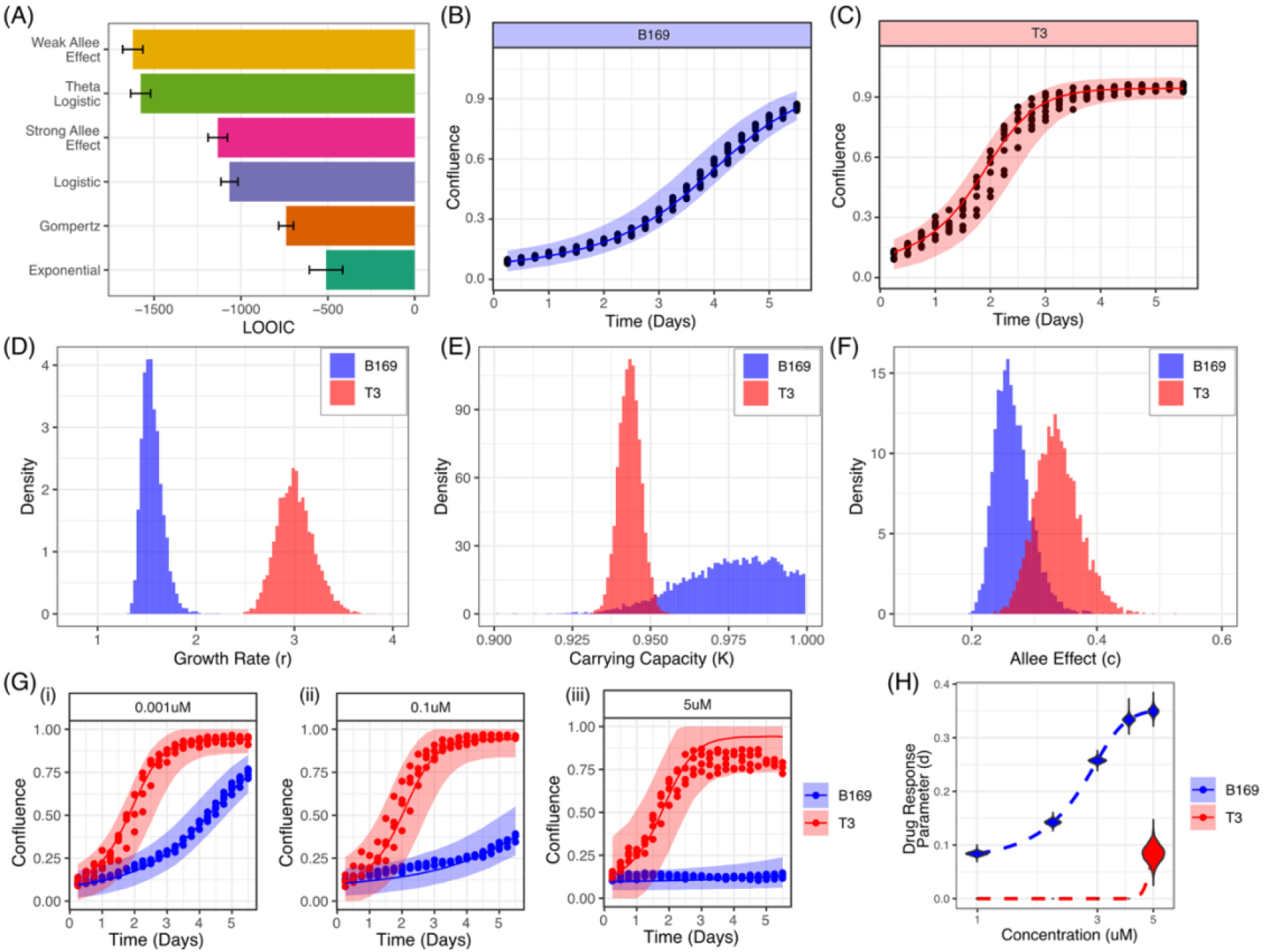
Parameterisation of mono-culture growth and therapeutic response dynamics B169 (blue) and T3 (red). **(A)** LOOIC values for common ODE models of growth supports the choice of a weak Allee effect model. **(B/C)** Allee effect model fitted to B169 and T3 with credible interval shaded. **(D)** Posterior distribution of fitted growth parameter, *r.* **(E)** Posterior distribution of fitted carrying capacity parameter, *K.* **(F)** Posterior distribution of fitted weak Allee effect parameter, *c.* **(G)** Fitted model for Trametinib response of B169 and T3 using a log-response with weak Allee effect model. **(H)** Violin plot over a range of concentrations of the posterior distribution for the drug response rate, *d*.

Upon deeper inspection of the weak Allee effect model, we can reproduce the dynamics of growth in B169 and T3, which are characterised by a sigmoidal pattern of growth (Figure 1B and 1C). Comparing this to a range of other models, we can see that the weak Allee effect model is able to closely follow both the early and late growth dynamics across all replicates. (Supplementary Figure 1)

The weak-Allee effect model consists of three parameters; the carrying capacity, growth rate and Allee effect threshold. The growth rate we observed is substantially higher for T3 (3.01 /day) compared to B169 (1.56 /day) (Figure 1D). The carrying capacity, which is the maximum size a population can attain, is comparable between the two lines, although B169 reaches a slightly higher maximal confluence (0.98 in B169 and 0.94 in T3) (Figure 1E). Finally, the Allee effect threshold is present for both lines, and is comparable between the two, although T3 has a slightly higher Allee effect threshold (0.34 in T3 compared to 0.27 in B169) (Figure 1F).

A visual inspection of the two lines highlights contrasting morphological qualities. B169 grows in dense clusters (Figure 1Gi) whilst T3 demonstrates growth dynamics that are substantially more diffuse (Figure 1Gii). This visually confirms our observations, as B169 displaying a slower growth rate in confluency, especially in the early dynamics.

As we previously mentioned, a defining distinction between these two lines is their response to trametinib. Trametinib potently inhibits B169 growth, whilst T3 shows little response [33]. We can introduce therapeutic intervention into the Allee effect growth model with a subtle modification:

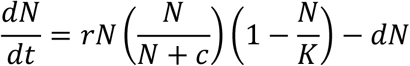

where an additional term introduced representing drug response is denoted as d. Fitting this model to the growth of parental and resistant monoculture under trametinib exposure from 0.001μM to 5μM, we can observe that the modified model is able to closely reproduce the dynamics of growth for both T3 and B169 under various concentrations of trametinib (Figure 1H). The posterior distribution of the response parameter (d) summarises the sensitivity to trametinib which confirms B169 is considerably more sensitive (Figure 1I). From the distribution of drug response parameters, we can see that the T3 line demonstrates no response at trametinib concentrations of up to 1uM and minimal response at 5uM.

### Detecting density-dependent interactions present within co-cultures

Interactions between distinct populations can be experimentally modelled and detected through the use of co-culture conditions [18,20]. Within our experimental set up, we have three distinct mixtures (25% B169-75% T3, 50% B169-50% T3 and 75% B169-25% T3). All cultures were seeded with the same number of total cells seeded, so a 50:50 co-culture would contain half as many B169 cells as the B169 monoculture, with the remainder of cells belonging to the resistant T3 population.

The Allee effect interaction predominantly affects the early growth dynamics present in these co-cultures, and thus looking closer at these dynamics may illuminate the presence of interactions. To achieve this, we fit an exponential growth model to the confluence values in the first 72 hours of each individual population in each co-culture and monoculture.

In the absence of a positive interaction, we would expect the growth rate of a population to decrease as the seeding density decreases. At lower ratios we will have fewer cells occupying a smaller collective area, therefore we expect a reduction in growth, which coupled with the presence of another directly competing population, would lead to a decline in growth rate. In general, competition decreases the growth rate of a population which has been demonstrated in several studies. [31,32,34]

On the other hand, a positive interaction between two distinct populations could either promote faster proliferation or diffusion, and this would be observed as a faster or similar growth in confluence in a mixture compared to in isolation. [18] This is beneficial for a population, since even though there are fewer cells occupying a smaller area initially, the presence of another population can promote growth in confluence at a faster rate than seeding a larger number of starting cells in a monoculture.

In the case of the T3 line, we can see that a reduction of the seeding ratio from a monoculture (seeding 100% Parental) to co-cultures, there is a reduction in the confluence after 3 days (Figure 2A). Since there are fewer T3 cells that occupy a smaller area coupled with the additional competition from the presence of B169 cells, our expectation of decreasing growth rate with decreasing seeding density is supported by the data (Figure 2B). Looking at images, we can see that there are simply fewer cells present, confirming growth is unsurprisingly limited by seeding at lower densities (Figure 2C).

**Figure 2:**
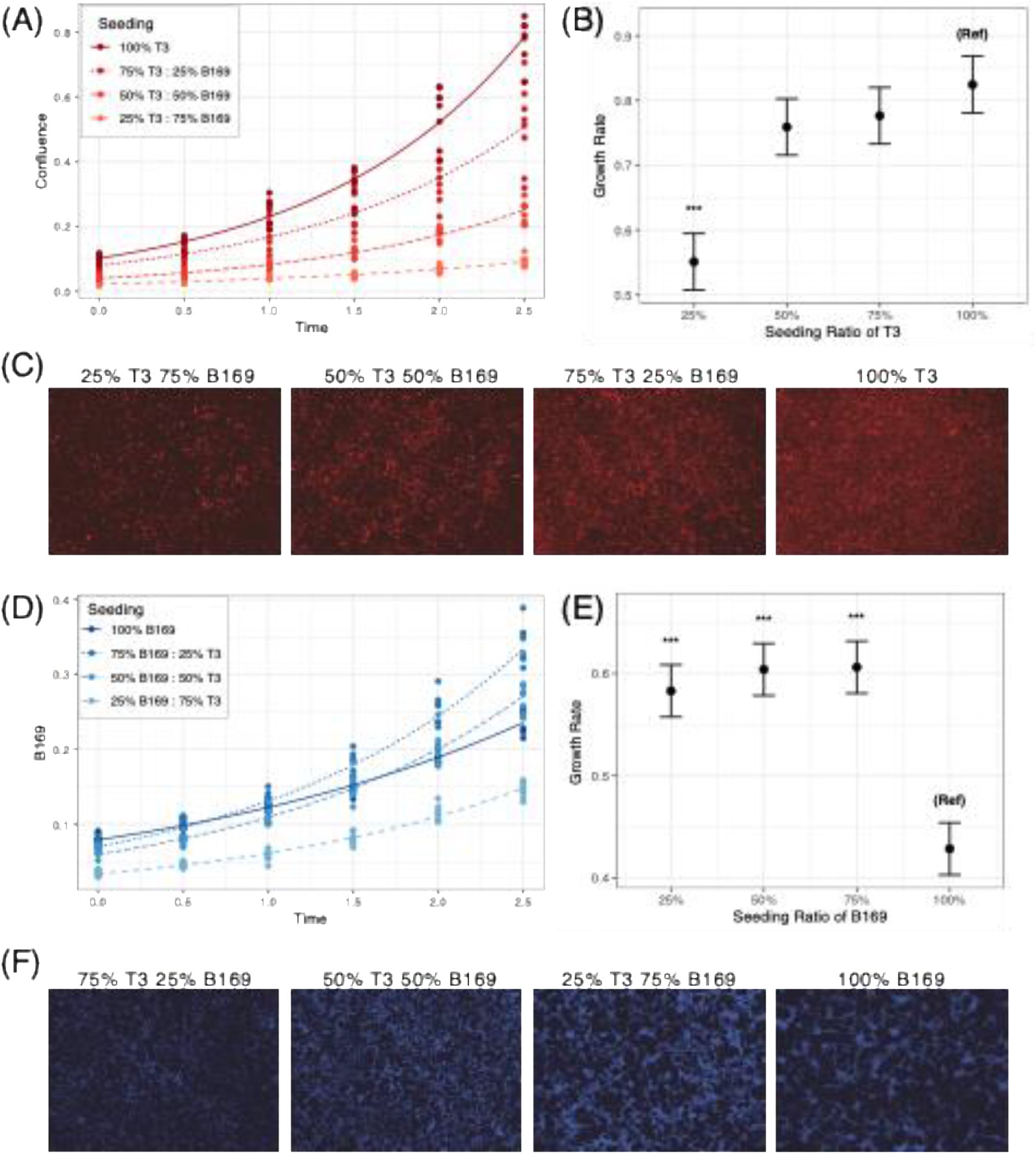
Early co-culture dynamics demonstrate an interaction enhancing the growth phenotype of B169. **(A)** 95% confidence interval of the fitted exponential growth parameter fitted to first 60 hours of growth for T3. Co-culture conditions display lesser growth in confluence. **(B)** Fitted parameter summary of exponential growth rate demonstrates slower growth as seeding density decreases for T3. **(C)** Images of red fluorescence channel from cultures at 60 hours supports the slower growth in confluence. **(D)** Exponential growth model fitted to first 60 hours of growth for B169. Co-culture conditions display higher or comparable growth in confluence despite lower initial confluence. **(E)** 95% confidence interval of the fitted exponential growth parameter fitted to first 60 hours of growth for B169. **(F)** Images of green fluorescence channel from cultures at 60 hours supports the faster growth in confluence of B169 in co-cultures.

Interestingly, this relationship is not observed in the case of the B169 line. When seeded at 50% or 75%, B169 line reaches a higher confluence than in monoculture, despite fewer starting cells (Figure 2D). The corresponding fitted exponential growth rate of B169 is significantly higher (p < 0.001) in all co-culture conditions (Figure 2E). Finally, we can see that this inference is supported by the experimental images, where B169 cells display larger confluence when seeded at 50% and 75% (Figure 2F). We also see a shift in the phenotype, from growing sparsely in clusters to displaying much more diffuse pattern of growth (Figure 2F).

### Quantifying co-culture interactions

We have demonstrated that the early-growth dynamics clearly differ, however, to quantify the observed interaction we require a system that is able to model the growth of both lines together. To achieve this, we will utilise a competitive Lotka-Volterra model, which is a frequently used model within cancer research. [31, 35] However, to incorporate the Allee effect we introduce an extension to this model by allowing interactions to modulate the Allee effect in addition to the existing Lotka-Volterra model. This leads to the following system of ODEs:

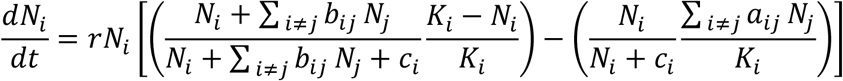

where the subscript denotes a distinct species of cell. As previously, N represents the population confluence, r represents the growth rate, K represents carrying capacity and c represents the Allee effect. The introduction of two terms; a_ij_ represents the Lotka-Volterra interaction and b_ij_ represents the introduced Allee effect interaction imparted on species i by species j. This system can be generalised to N distinct species; in this study we utilised a system with two distinct species.

Using simulations, we can demonstrate which of the two interaction sources can produce a change in the early-stage co-culture growth dynamics akin to those we observed previously. These simulations vary the strength of either the Lotka-Volterra interaction or Allee effect interaction, while keeping the other fixed. The Lotka-Volterra interactions parameters are contained within the ‘occupancy term’ – a term we use to describe the ‘fullness’ of an environment. These interactions typically have a larger effect when the total occupancy is greater, naturally this will be less impactful at early dynamics (Figure 3A i) compared to those at later stages when the system is closer to carrying capacity (Figure 3A ii). The Allee effect interactions, however, are more impactful at lower densities (Figure 3A iii) and thus more prominently affect the initial growth dynamics, with the Allee effect diminishing as the population density increases and the effect of interaction subsiding (Figure 3A iv).

**Figure 3:**
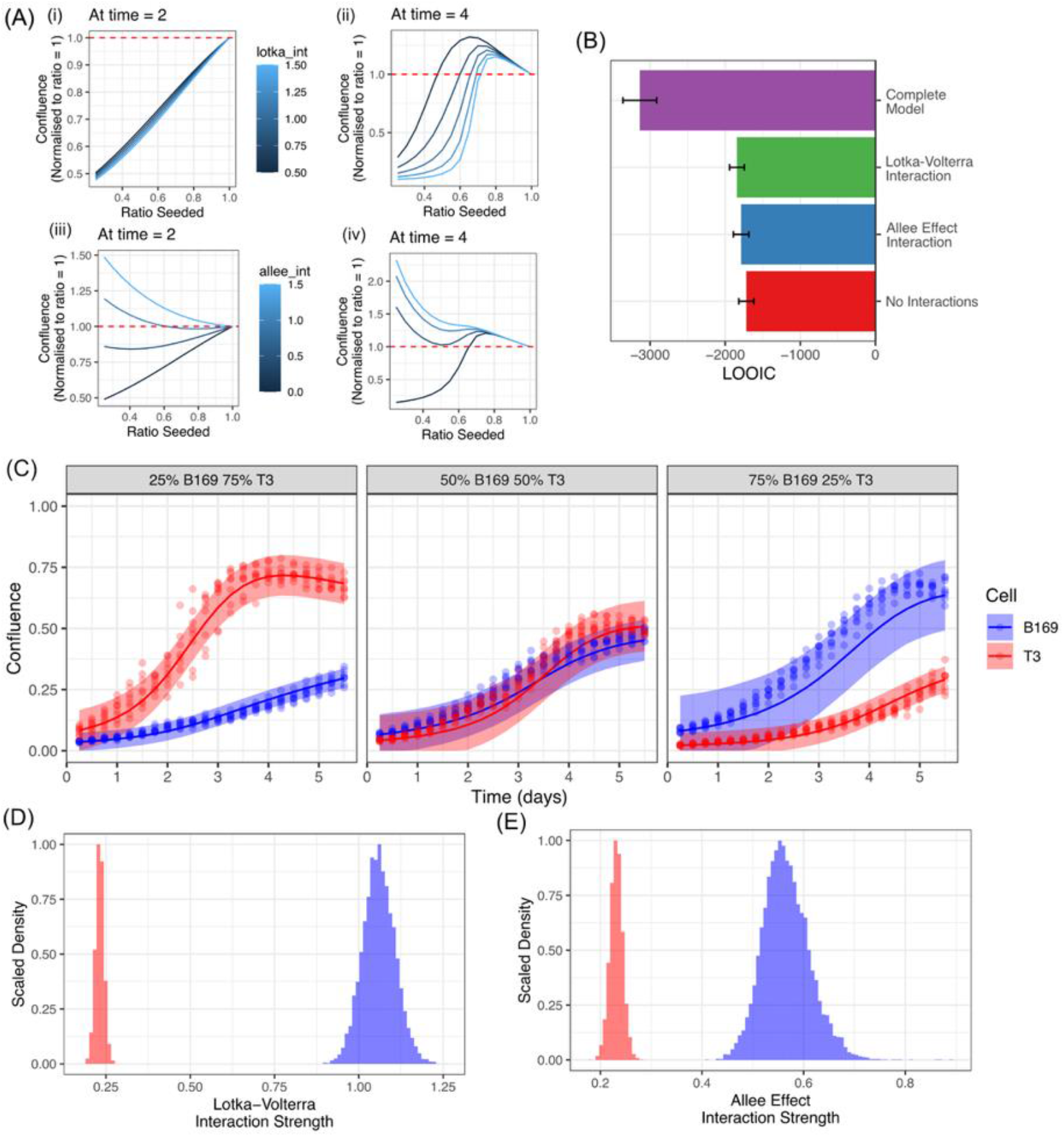
Inference of co-culture dynamics demonstrates the presence of an interaction impacting the Allee effect. **(A)** Simulations to understand the impact of Lotka-Volterra and Allee effect interactions on the growth rate normalised by confluence. **i/ii)** Lotka-Volterra interactions shows increased confluence in co-cultures which is more pronounced at later stages of growth. **iii/iv)** Allee effect interactions impact the earlier dynamics with this increase in confluence becoming less impactful at later stages of growth. **(B)** LOOIC of permutations of interaction models demonstrates a model with both Allee effect and Lotka-Volterra interactions best fit the data. **(C)** Fitted interaction model with both Allee effect and Lotka-Volterra interactions with a single set of parameters demonstrates reasonable alignment with the data. **(D)** Posterior distribution of Allee effect interaction demonstrates a substantially higher impact of interactions on B169. **(E)** Posterior distribution of Lotka-Volterra interactions demonstrates B169 is impaired by competition to a higher degree than T3.

Applying these models to infer interactions, we explored permutations of variants with interactions included or excluded. We leverage the previously parameterised monoculture phenotype of these lines and integrate this into the extended interaction model. The best fitted model is one with both Allee and Lotka-Volterra interactions, as demonstrated by this model possessing the lowest LOOIC (Figure 3B). This model is best able to track the dynamics of B169 and T3 across all three co-cultures, being able to reproduce the inflection points and possessing the narrowest credible intervals of the inflection (Figure 3E). A model with solely Lotka-Volterra interactions is unable to capture the dynamics (Supplementary Figure 2A), whilst a model with only Allee effect interactions has a less certain fit (Supplementary Figure 2B).

Looking at the posterior distributions of the final chosen model highlights the nature of the interaction we observe. Aligning with our prior observations where the sensitive line grows substantially faster in a co-culture than in isolation, we can see that the Allee effect interaction experienced by B169 is substantially higher than that experienced by T3 (Figure 3F). We also observe that the resistant line is relatively unperturbed by the Lotka-Volterra interaction, with a value considerably below 1, while the sensitive line experiences substantially more competition comparatively (Figure 3G). The implication of this interaction is that at low densities, the sensitive population derives an advantage growing and occupying a wider area than it would have in isolation, however, in the longer-term dynamics the resistant line would dominate our experimental environment.

### Therapeutic Intervention

A natural extension to our experimental system, with trametinib sensitive and resistant populations, is to understand how the interactions we have observed interplay with the introduction of therapy. We have already demonstrated that trametinib is not tolerated by the sensitive line, with the resistant line showing minimal response.

Looking at the relative confluence of sensitive cells in the first 3 days of drug exposure, the growth rate of these cells demonstrates the presence of an impact from a positive interaction in the two lowest concentrations 0.01uM and 0.001uM. However, once the concentration of trametinib is increased to 0.1uM the interaction disappears, in this condition the growth is considerably inhibited (Supplementary Figure 3). This may be due to the interaction no longer being able to compensate for the competition present.

We propose a modification of the existing co-culture model to include therapeutic response, with interactions that can be modified by the inclusion of therapy. Given the knowledge that the resistant line does not respond to therapy, we focus on understanding the impact on the parental line. In a co-culture setting, we could expect not only the growth parameters to be perturbed but also the interaction parameters we previously identified.

The model we fit is now:

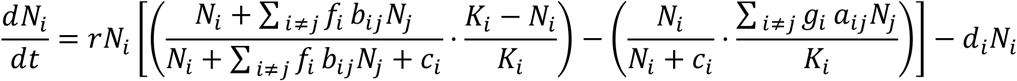

Iterating through various permutations, we observe the best model to involve the Lotka-Volterra and Allee effect interactions being perturbed in line with therapeutic response, demonstrated by this model possessing the lowest LOOIC (Figure 4A).

**Figure 4:**
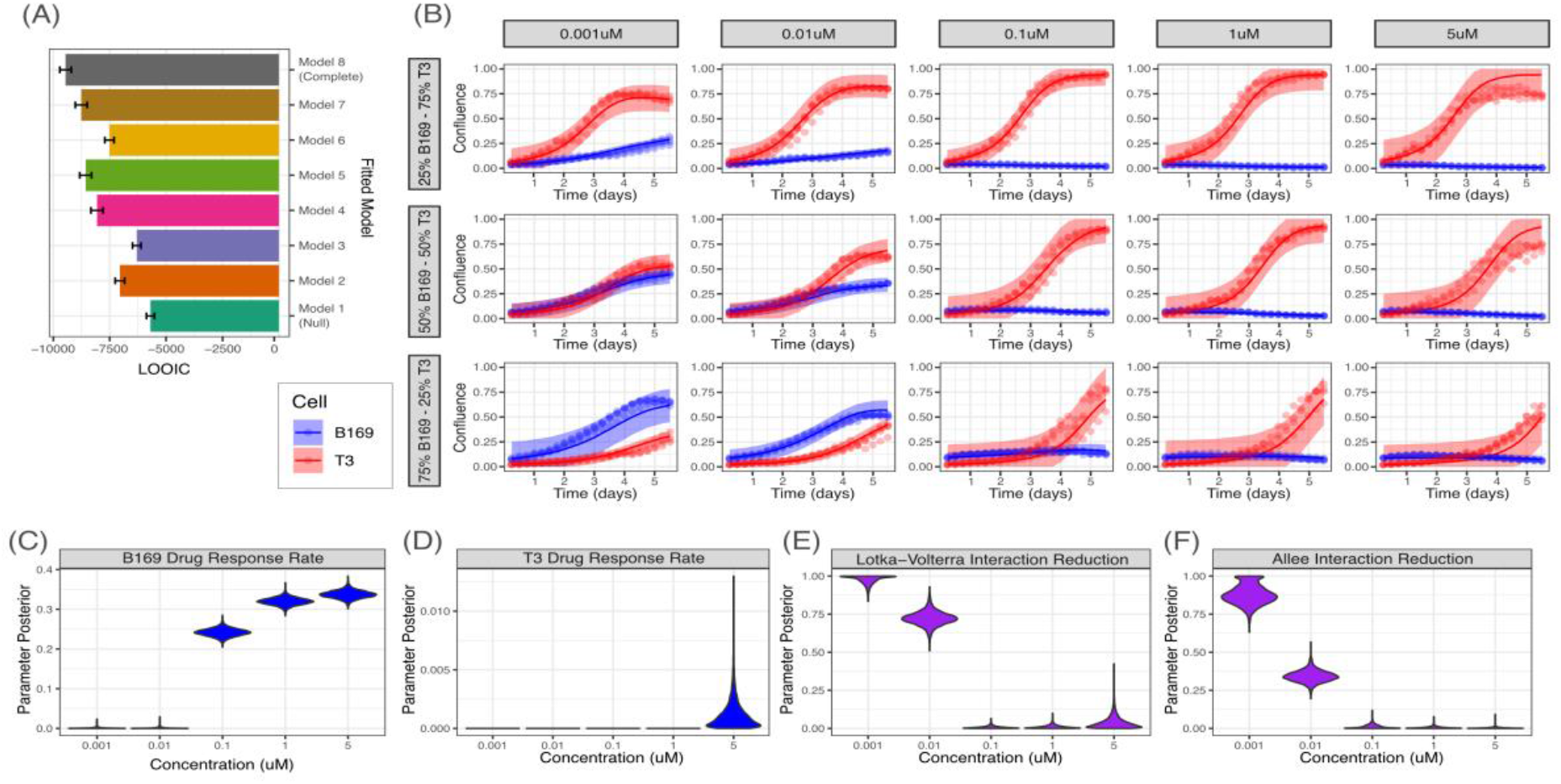
Therapeutic intervention with trametinib perturbs observed interactions eliminating the observed cooperativity. (A) LOOIC of explored therapeutic response models demonstrates perturbation of interaction in line with the therapeutic target best aligns with the data. (B) Fitted model for B169 and T3 co-cultures at trametinib concentrations ranging from 0.001uM to 5uM. (C) Posterior distribution of drug response parameter aligns with mono-culture therapeutic response for B169 except for lower concentrations. (D) Posterior distribution of drug response parameter aligns with mono-culture therapeutic response for T3. (E) Posterior distribution of Lotka-Volterra interaction parameter demonstrates a dose-dependent reduction interaction for T3, highlights a lesser impact of competition. (F) Posterior distribution of Allee effect interaction parameter demonstrates a dose-dependent reduction in the Allee effect interaction received by B169, highlighting the elimination of the interaction.

The fitted model can capture the dynamics of both B169 and T3 in all three co-cultures across the range of trametinib concentrations (Figure 4B).

Looking at the posterior distribution of fitted parameters, we can see that the drug response rate of B169 is lower for concentrations 0.001uM and 0.01uM compared to a monoculture response (Figure 4C). When looking at this parameter for T3 we observe minimal change and T3 still demonstrates resistance to trametinib (Figure 4D).

In the selected model, the Lotka-Volterra interaction (representing the competition between lines) is reduced for the resistant line as trametinib concentration is raised (Figure 4E). Intuitively, this aligns with the response we observed as sensitive cells are impacted by trametinib therapy. Similarly, we can see the Allee effect interaction received by the sensitive cells from resistant cells is also reduced (Figure 4F); here the sensitive cells could be seen as less able to receive an interaction when treatment is applied. Together, these finding suggest that an interaction confers a growth advantage in co-cultures that persists at low trametinib concentrations and is lost at higher doses.

### Interactions may modulate the optimal sequencing of therapeutic interventions

To understand the impact these interactions could potentially have in a two-drug system, we created a toy model. Here the two modelled cell lines have their growth rates (r = 1.5), Allee effect (c = 0.25), carrying capacity (K = 0.95) and Lotka-Volterra interaction (b_ij_ = 1) equalised. The only factor varied is the Allee effect interaction (a_AB_ = 1, a_BA_ = 0), with only population A receiving the interaction. We set population A as sensitive to Drug 1 (trametinib) and population B sensitive to Drug 2 (dasatinib), with each sensitive population having resistance to the other drug.

We can first see that a single continuous therapy would lead to an uncontrolled increase in the confluence of the untargeted population (Figure 5A). This implies the failure of a simple single drug regime. When trametinib is applied the trametinib resistant population dominates the culture (Figure 5A i), and when dasatinib is applied the dasatinib resistant population is selected for (Figure 5A ii).

**Figure 5:**
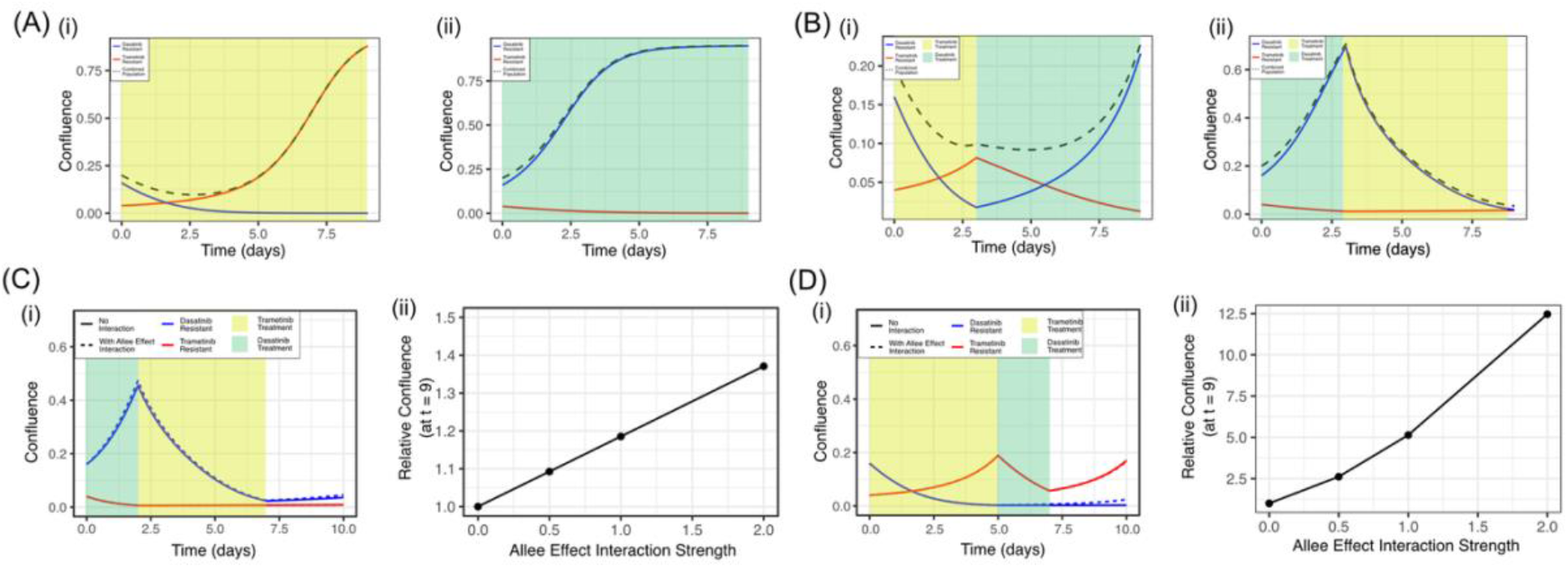
Simulated exploration of a two-drug paradigm with the presence of Allee effect interactions demonstrates the potential to target interactions to obtain better control of tumour response. (A) A single drug protocol leads to the failure of intervention as the resistant population dominates. (B) A two-drug sequential system i) Trametinib followed by dasatinib ii) Dasatinib followed by trametinib. (C) Scheduling dasatinib prior to trametinib followed by a recovery period leads to similar outcomes in the presence and absence of Allee effect interactions. i) Overall confluence of subpopulations with and without and interaction. ii) Relative confluence of the subpopulation that benefits from the interaction at t=9 after 3 days of treatment being removed. (D) Scheduling trametinib prior to dasatinib followed by a recovery period leads to deviations in the outcomes under the presence and absence of Allee effect interactions. i) Overall confluence of subpopulations with and without and interaction. ii) Relative confluence of the subpopulation that benefits from the interaction at t=9 after 3 days of treatment being removed.

With multiple options of therapies, conventional wisdom would advise targeting of the largest population first. In this case we start by targeting the larger population A with trametinib followed by dasatinib, with the result that the incomplete removal of the trametinib sensitive population leads to its expansion when treatment is lifted (Figure 5B i). Interestingly, when dasatinib is applied first with identical lengths of treatment, we can see that the overall population at the end of treatment sequencing is considerably lower (Figure 5B ii). This demonstrates that there is merit in not simply targeting the largest population first but considering the diversity of the population.

To understand the potential mechanism that may underlie this observation, we can look at scaling the Allee effect interaction strength from absent (at 0) and strong (at 2). In the case where the population is maintained under stricter control, dasatinib followed by trametinib treatment, we can see that there is minimal difference between the case with and without interactions (Figure 5C i). There is a positive relationship between the interaction strength and relative confluence of the interaction-receiving dasatinib resistant population (Figure 5C ii).

The striking observation is when this treatment protocol is flipped so that the dasatinib resistant population is targeted first. Here we see a clear distinction between the case of interactions and no interactions, with higher confluence when interactions are present (Figure 5D i). Most importantly, the interaction strength has a considerably stronger relationship with the relative confluence of the interaction receiving population (Figure 5D ii).

Under this system, targeting the largest population when it can receive interactions from a smaller underlying population leads to less strict control of tumour growth. Instead, targeting the interaction-providing population first appears to prime the system to respond better to treatment. These observations demonstrate that there is utility in devising therapeutic strategies with the view of the targeting of interactions between subpopulations and not just the individual subpopulations.

## Discussion

This study presents a distinct extension to mathematical models designed specifically to interpret interactions between distinct populations within a competitive environment. Here, we expanded upon the commonly used, Lotka-Volterra competition model, by introducing a weak Allee effect and allowing for an interaction to perturb its impact. Both the Lotka-Volterra and weak Allee effects are widely studied within biological systems, particularly in cancer. For example, Zhang et al. leveraged a competitive Lotka-Volterra model to understand the principles of adaptive therapy [31] Gallagher et al. developed a framework of using Lotka-Volterra models in clinical settings to manage disease longer term. [35] The Allee effect is less commonly studied in the literature as it is a phenomenon primarily restricted to smaller populations, nonetheless, Gerlee et al. explored the role of autocrine signalling in the emergence of an Allee effect within cancer cell populations [23]. The modulation of an Allee effect through community interactions has also been proposed by Hersey et al [36].

We conducted a coupled experimental and computational investigation to quantify the presence of interactions that impart a growth advantage to cells in a heterogenous environment. We showed this interaction could be quantified and understood through the lens of an Allee effect, and proposed a mathematical formulation of this phenomenon, building upon existing competitive growth models. We utilised confluence as a key metric to track the growth dynamics of the individual populations, due to its relevance within paediatric high-grade gliomas, specifically DMG. These tumours typically develop aggressively and in a diffuse pattern, and thus confluence allows for the incorporation of both cell proliferation and cell dispersion into a single metric. However, a limitation with this approach is the inability to dissect between proliferation and dispersion.

First, we demonstrated that a weak Allee effect model represents the best description for the growth dynamics present in these cultures. Using this representation, we were able to parameterise the growth dynamics in isolation. This parameterised description of population is used to condition subsequent inference on both co-culture and therapeutic dynamics.

Within co-cultures we noticed a pronounced increase in the confluence of only one subpopulation, B169, when compared to the phenotype in isolation. We were able to attribute this visually to a distinct change in the modality of growth, with co-cultured B169 cells displaying a more dispersed phenotype. This increase in confluence was observed from an early stage, which we demonstrated to be incompatible with a typical Lotka-Volterra interaction model with and without Allee effects. Given the Allee effect is a community-driven phenomenon, we demonstrated that an interaction that alleviates the Allee effect can recapitulate the initial growth dynamics. This produces the key model introduced within this study, which we validate produces the best fit from a panel of alternative models. This enables for the quantification of the interaction parameters, the result of which supports the observations of early growth increases, with B169 receiving a credible contribution to its Allee effect from T3.

To further demonstrate the strength of these findings, we explored therapeutic dynamics as B169 and T3 have differential responses to trametinib. From this exploration, we were able to parameterise the response to treatment supporting the differential sensitivity. This highlighted that the interactions present were also perturbed by treatment and at low concentrations they lessened the burden of treatment. Extending this with simulations, we were able to show the presence of interactions to have considerable implications for treatment sequencing, favouring targeting of the interaction over simply treating the largest population first. In fact, there was stricter control achieved by eliminating a population that provides an interaction even if it was present at a lower abundance. This aligns with an active area of exploration, where combination therapies with trametinib have been explored in PDHGG that have produce improvements in patient overall survival. [37,38]

There are a multitude of avenues for further expansion of both the mathematical models and biological implications. A primary goal would be to understand the biological factors that drive the interaction between B169 and T3. We observed both a change in growth dynamics as well as a markedly different phenotype for B169, with a more dispersed presentation in co-cultures. Potential explanations of this switch could be through non-cell autonomous interactions that activate signalling pathways, such as those involved in a mesenchymal transition. If these interactions can be biologically validated, it would be possible to utilise existing or develop novel therapeutics to target them. It remains to be seen whether these Allee effect interactions are present within a tumour *in situ*. The nature of these interactions is that they occur at lower densities, however, early-stage tumours are seldom detected and rarely studied in-depth.

A limitation in this study is the lack of the presence of cells that make up the non-tumour microenvironment, which have been shown to provide an advantage to malignant cell growth [21,23]. Since the Allee effect is presented in small populations that lack the community effects to grow optimally, there is the prospect that cooperativity between tumour and non-tumour cells may enable the early development of tumours and support their spread. While this limitation exists, it also represents a distinct opportunity to understand whether and how the tumour microenvironment sustains the conditions for optimal tumour growth. The ability to detect these non-tumour-to-tumour interactions would represent a crucial tool in dissembling the complexity in the tumour microenvironment. This study provides both the computational methodology and an experimental approach to detect and quantify such interactions.

To summarise, through the generation of mathematical models seeking to understand the dynamics of interactions that alleviate the Allee effect, this study highlighted their presence within patient-derived DMG models. These interactions were shown to enhance the diffuse pattern of growth, maintaining relevance in a therapeutic environment. This raises the implication that microenvironmental cooperation, within tumours and potentially with their surroundings, represent a clinically exploitable avenue for tumour growth, dispersion and therapeutic escape.

## Materials and Methods

### In-vitro cell culture

DIPG biopsy tissue was shipped to our laboratory in Hibernate A transport media (ThermoFisher Scientific, A12475-01) at room temperature or minced with a sterile scalpel blade in DMEM/F12 (Life Technologies, 11320-074) supplemented with 0.2% BSA (Sigma-Aldrich, A1595) and 10% DMSO (Sigma-Aldrich, D2650), frozen at −80°C and sent on dry ice (Supplementary Table S1). Minced tissue was digested using Liberase DL. Supernatant was removed and the tissue/cells were resuspended in stem cell media and continuously pipetted to ensure dissociation before transfer to culture flasks to grow either attached on a laminin substrate (Merck Millipore,

CC095) and/or in suspension as neurospheres in ultra-low attachment flasks (Sigma-Aldrich, CLS3815). The cells were grown at 37°C, 5% CO2 in stem cell media consisting of DMEM/F12 (Life Technologies, 11330-038), Neurobasal-A Medium (Life Technologies, 10888-022), HEPES Buffer Solution 1 mol/L (Life Technologies, 15630-080), MEM Sodium Pyruvate Solution 100 nmol/L (Life Technologies, 11360-070), MEM Non-Essential Amino Acids Solution 10 mmol/L (Life Technologies, 11140-050), and Glutamax-I Supplement (Life Technologies, 35050-061). The media were supplemented with B-27 Supplement (Life Technologies, 12587-010), 20 ng/mL recombinant Human-EGF (2B Scientific LTD, 100-26), 20 ng/mL recombinant Human-FGF (2B Scientific LTD, 100-146), 20 ng/mL recombinant Human-PDGF-AA (2B Scientific LTD, 100-16), 20 ng/mL recombinant Human-PDGF-BB (2B Scientific LTD, 100-18), and 2 μg/mL Heparin Solution (Stem Cell Technologies, 07980).

### Co-culture growth assays

In 96-well laminin coated plates, 1000 cells per well were seeded from a stock consisting of a monoculture of ICR-B169-Parental or ICR-B169-T3, as well as three ratios consisting of 750, 500 and 250 cells of ICR-B169-Parental, and 250, 500, and 750 cells of ICR-B169-T3 respectively. Cells were seeded with 100uL of stem cell growth medium and allowed to attach for 24 hours. After attachment 100uL of medium was added with either DMSO, trametinib or dastanib to achieve the desired concentrations of 0.001uM to 5uM. Cells were then placed in an IncuCyte S3 live-cell imaging system in a humidity and CO2 controlled incubator for 6 days.

### Cell labelling

To generate stable fluorescent labelled cells, cells were engineered using the PiggyBac transposon system to constitutively express eGFP (B169-Parental) and mCherry (B169-T3). Plasmids were propagated in Subcloning Efficiency DH5α competent cells and isolated using a QIAprep Spin Miniprep Kit (Qiagen).

Transfection was performed by nucleofection using the Human Stem Cell Nucleofector Kit 2 (Lonza, VPH-5022) on a Lonza Nucleofector device.

Briefly, 8×105 cells per reaction were harvested, resuspended in 100 μL of supplemented Nucleofector Solution 2, and combined with recommended amounts of transposon donor and transposase helper plasmids (1–5 μg total DNA). Samples were pulsed using program A-23, then immediately placed in cell culture medium and plated into T-25 culture flasks. Cells were allowed to recover to achieve >50% confluence and after confirmation of fluorescence signal, cells were sorted using FACS to isolate positively labelled cells. Following this, labelled cells were further expanded and confirmed to be stably labelled.

### Cell Imaging

Cells were imaged in the live-cell imaging system, Satorius IncuCyte S3. Seeded cells were allowed 12 hours to acclimate and the imaged at 6-hour intervals using a 4x magnification, with red and green-fluorescent channels to detect mCherry and eGFP labels respectively.

### Monoculture ODE model

We use a logistic growth model with weak Allee effect defined as follows:

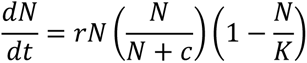

where *r* represents the growth rate, N represents the confluence of the population, *K* represents the carrying capacity (the maximum confluence the population can attain) and *c* represents the weak Allee effect strength.

### Co-culture ODE models

The final co-culture model utilised involves a modified Lotka-Volterra model with Allee effects where interactions are also able to perturb the Allee effect. This is defined as follows:

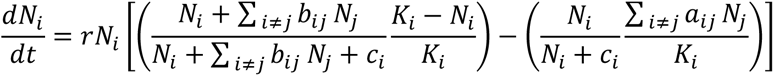

where the subscript denotes the population and added parameters b_ij_ the Lotka-Volterra interaction population i receives from population j, a_ij_ the Allee effect interaction population i receives from population j.

### Therapeutic response model

The therapeutic response involves a subtraction from the growth rate in line with the strength of treatment. The response is represented with the parameter *d* which is added to the monoculture and co-culture models as follows:

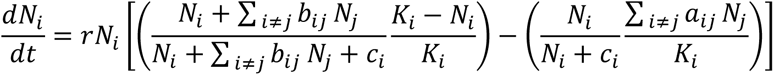

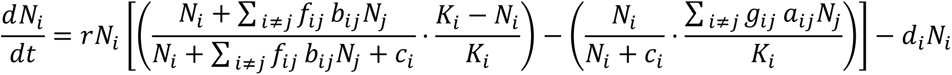

in the co-culture model, the parameters *f* and *g* are multiplicative perturbations of the interactions which allows for treatment to also impact the interaction. This parameter is multiplicative so it can be absorbed into the interaction term for the purpose of inference.

### Bayesian Inference

Bayesian Inference was performed in R using the rstan library. [38] Using an ode45 solver, generated quantities were fit to data with normally distribution residuals. Non-informative uniform priors were used for monoculture models, with limits established based on intuitive limits (such as limiting carrying capacity between 0 and 1).

Subsequent models that had parameters that were shared with simpler models, drawn from the posterior distribution of those simpler models. For example, a co-culture model sharing growth and carrying capacity parameters with a monoculture model. All analysis used 4 chains with 2000 iterations and 1000 warmup iterations. LOOIC (leave-one-out cross-validation information criteria) was calculated using the loo function from the loo library in R.

## Supplementary Figures

**Supplementary Figure 1:**
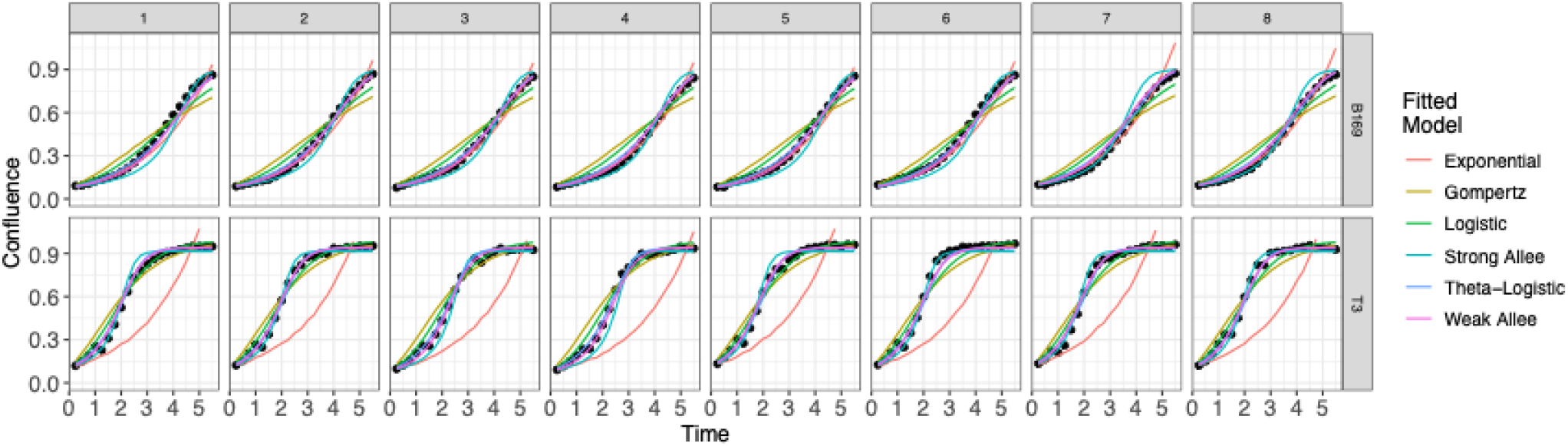
Best fit line from a range of mono-culture growth models.

**Supplementary Figure 2:**
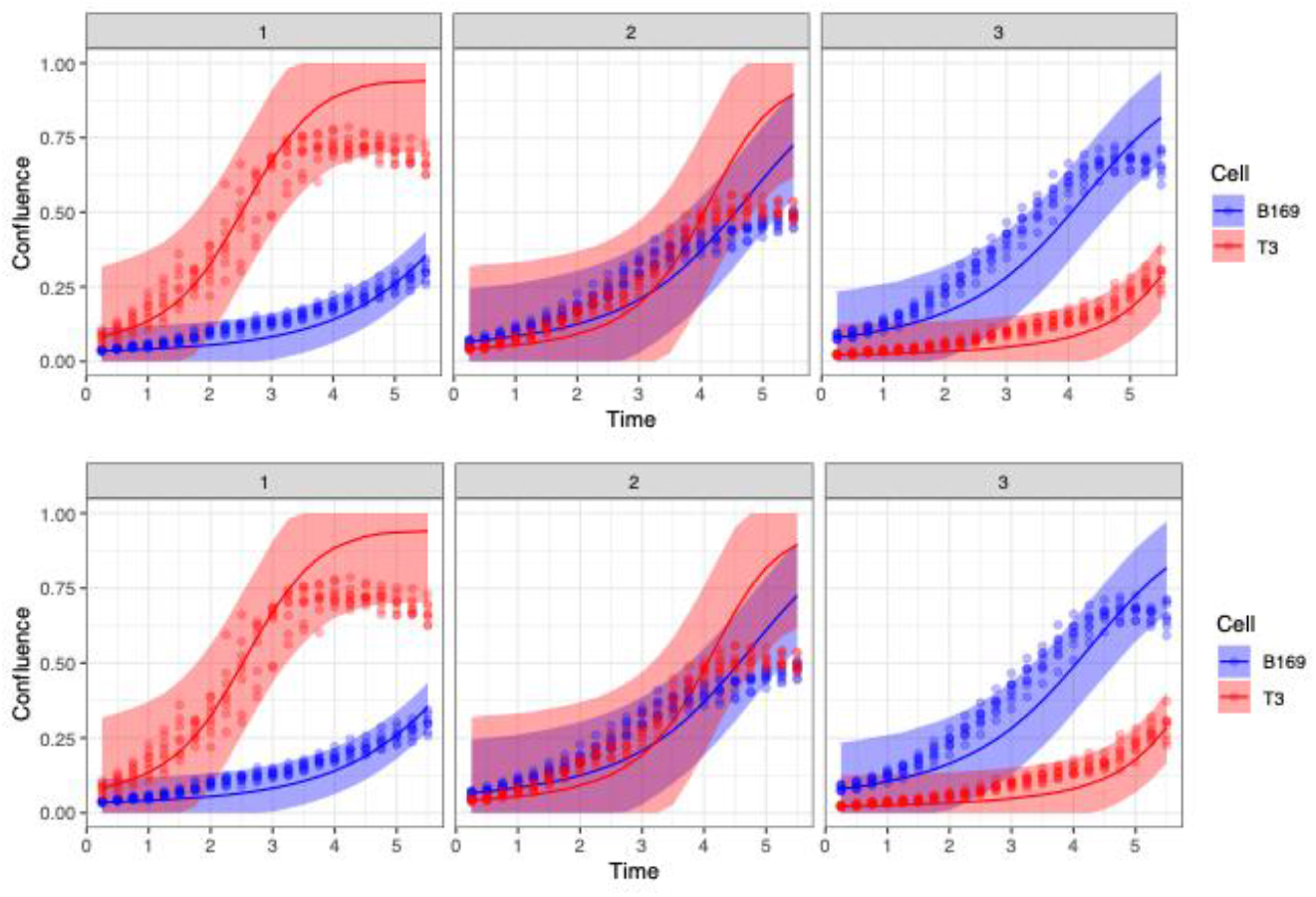
A) Resulting fit of a co-culture model fit with only Lotka-Volterra interactions **B)** Resulting fit of a co-culture model fit with only Allee effect interactions.

**Supplementary Figure 3:**
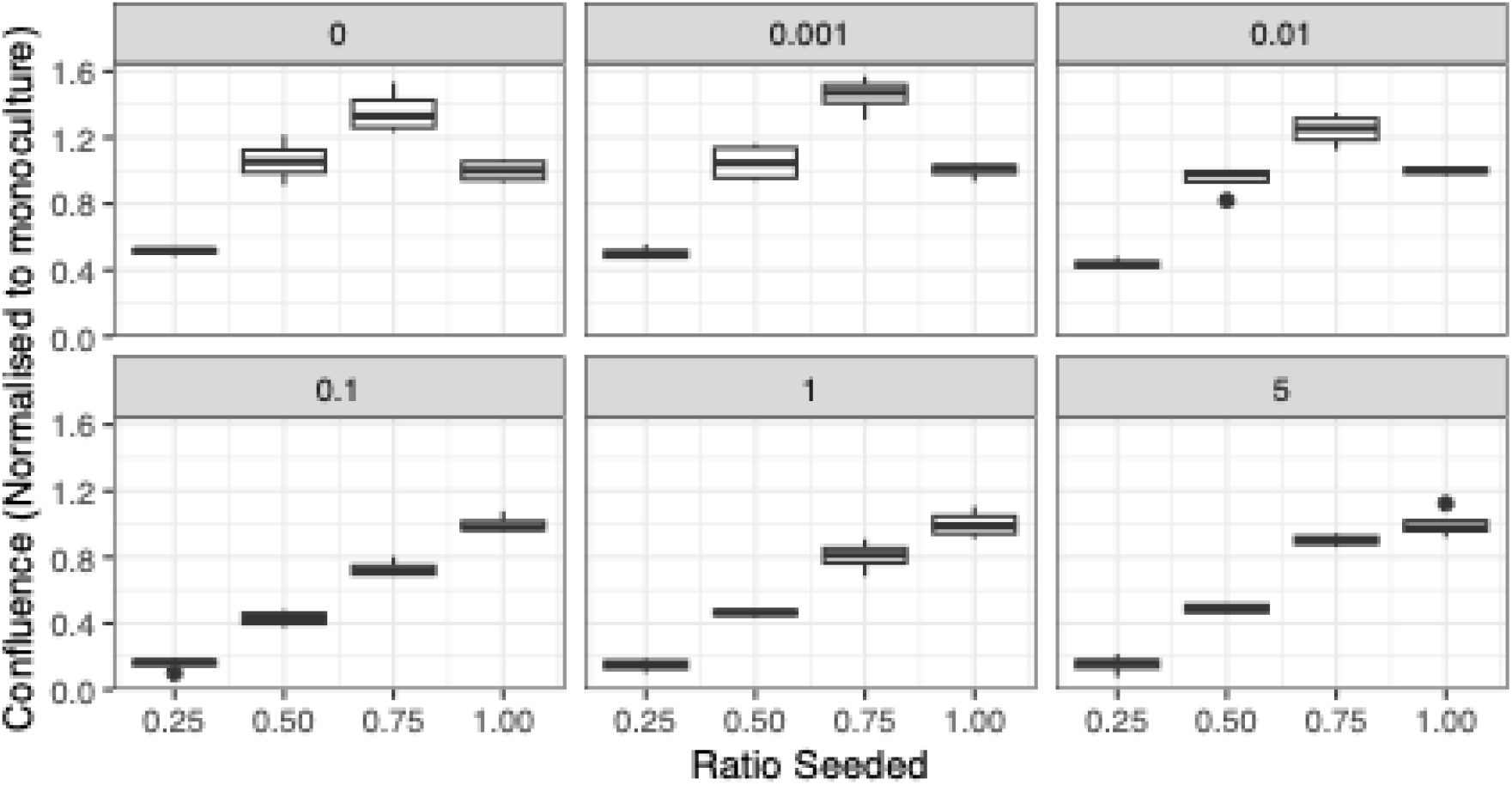
Confluence of B169 across monoculture and co-culture conditions normalised to monoculture (Ratio Seeded = 1) under trametinib therapy from 0uM to 5uM.

